# A game theoretic approach to deciphering the dynamics of amyloid-β aggregation along competing pathways

**DOI:** 10.1101/581629

**Authors:** Preetam Ghosh, Pratip Rana, Vijayaraghavan Rangachari, Jhinuk Saha, Edward Steen, Ashwin Vaidya

**Affiliations:** Department of Computer Science, Virginia Commonwealth University, Richmond, VA 23220; Department of Chemistry & Biochemistry, School of Mathematics and Natural Sciences, University of Southern Mississippi, Hattiesburg, MS 39406; Department of Mathematical Science, Montclair State University, Montclair, NJ 07043

## Abstract

Aggregation of amyloid *β* (A*β*) peptides is a significant event that underpins Alzheimer disease (AD). A*β* aggregates, especially the low-molecular weight oligomers, are the primary toxic agents in AD pathogenesis. Therefore, there is increasing interest in understanding their formation and behavior. In this paper, we use our previously established investigations on heterotypic interactions between A*β* and fatty acids (*FA*s) that adopt off-fibril formation pathway under the control of *FA* concentrations, to develop a mathematical framework in defining this complex mechanism. We bring forth the use of novel game theoretic framework based on the principles of Nash equilibria to define and simulate the competing on- and off-pathways of A*β* aggregation. Together with detailed simulations and biophysical experiments, our mathematical models define the dynamics involved in the mechanisms of A*β* aggregation in the presence of *FA*s to adopt multiple pathways. Specifically, our game theoretic model indicates that the emergence of off- or on-pathway aggregates are tightly controlled by a narrow set of rate constant parameters, and one could alter such parameters to populate a particular oligomeric species. These models agree with the detailed simulations and experimental data on using *FA* as a heterotypic partner to modulate temporal parameters. Predicting spatiotemporal landscape along competing pathways for a given heterotypic partner such as biological lipids is a first step towards simulating physiological scenarios in which the generation of specific conformeric strains of A*β* could be predicted. Such an approach could be profoundly significant in deciphering the biophysics of amyloid aggregation and oligomer generation, which is ubiquitously observed in many neurodegenerative diseases.

## 1. Introduction

Aggregation of the protein amyloid-β (Aβ) is one of the central processes in the etiology of Alzheimer disease (AD). Generated by the proteolytic processing of amyloid precurssor protein (APP), Aβ peptides (Aβ40 or Aβ42) spontaneously aggregate to form insoluble fibrils that deposit as senile plaques in the AD brain. In the aggregation pathway, the low molecular weight oligomers are the primary toxic species that are responsible for synaptic dysfunction and neuronal loss(Chromy et al., 2033; Cleary et al., 2005; O’Brien and Wong, 2011; Smith et al., 2006; Walsh et al., 2002). An increasing number of reports indicate that strucutral polymorphism and heterogenity within the aggregates could contribute to clinical phenotypes observed among AD patients (Selkoe, 2001). Therefore over the last decade, significant efforts are focused on understanding the biophysical and biochemical aspects of aggregation as well as the molecular understanding of the aggregates.

It is now well established that aggregation follows a nucleation-dependent, sigmoidal growth kinetics. Although the precise mechanism of nucleation continues to be heavily debated, substantial evidence suggests a rate-limiting mechanism for the formation of nucleus or nuclei (Buell et al., 2014; Cohen et al., 2013; Ghosh, Datta, Rangachari, 2012; Ghosh et al., 2010; Rangachari et al., 2007). Since the nucleation plays an important role in determining the morphology of the fibrils formed, the dynamics associated with reactions leading up to nucleation is critical for aggregation. Intrinsic disorder and amphipathic nature of monomeric Aβ makes it particularly sensitive to environmental factors and other interacting partners (Fraser, Darabie, McLaurin, 2001; Kirkitadze, Condron, Teplow, 2001; Kumar et al., 2011; Nichols et al., 2005; Wood et al., 1996), and thus, Aβ is known to adopt multiple pathways depending on the aggregation conditions. As a result, it is clear that oligomers may not be the obligate intermediates of fibril formation, and oligomers with disctinct conformations can also be formed along alternative aggregation pathways (off-pathways) (Bitan et al., 2003; Gellermann et al., 2008; Goldsbury et al., 2005; Liu et al., 2012; Necula et al., 2007; Rangachari et al., 2007). This is significant because such interactions, depending on the strucutre of the oligomer, determine the morpholgy of the aggregates formed and consequently the toxicity and possibly the phenotypes.

As competing aggregation pathways could result in distinct oligomer strains with pathological implications, it is imperative to gain understanding on how physiological interacting partners of Aβ affect its aggregation dynamics. Being generated from the membrane spanning domain of the amyloid precrsor protein (APP), Aβ displays almost synchronous and perpetual interaction with membrane lipids (Choo-Smith et al., 1997; Choo-Smith and Surewicz, 1997; Fletcher and Keire, 1997; Kakio et al., 2002; Kim, Yi and Ko, 2006; Williams, Day and Serpell, 2010; Yong et al., 2002). Therefore, amphipathic surfactants and lipids have a strong ability to modulate Aβ’s conformation to drive towards multiple aggregation pathways. Interfaces of lipids and fatty acids are of profound interest in physiological contexts as they are abundant in both cerebral vasculature and CSF (Carpentier and Hacquebard, 2006; Schlame et al., 1996). Previous reports from ours and other laboratories have established that phase transitions of surfactants and membrane lipids modulate Aβ aggregation pathways in a concentration dependent manner to generate aggregates via at least one alternative, off-pathway from the canonical fibril formation pathway (on-pathway) (Power and Powers, 2008; Rana et al., 2017; Kumar et al., 2011; Gellerman et al., 2008; Rangachari et al., 2007; Gorman et al., 2003; Kumar et al., 2012; Solfrizzi et al., 2011). Specifically, low-molecular weight oligomers were generated in the presence of fatty acid near and above their respective critical micelle concentrations (CMC) (pseudo-micellar and micellar, respectively) and not below CMC (non-micellar), which augmented the fibril formation, on-pathway (Kumar et al., 2011, Rana et al., 2017; Rangachari et al., 2018). More importanty, the oligomers generated were deemed to be formed along the off-pathway based on their conformation and half-lives (Rana et al., 2017; Kumar et al., 2012).

The adoption of multiple aggregation pathways by Aβ, along with the influence of heterotypic interactions in modulating them posit the question of what spatiotemporal parameters guide the modulatory dynamics, and whether one could simulate the temporal emergence and dissappeance of aggregates as a function of heterotypic Aβ interactions. In this work, we have approached to answer these questions using the well-establsihed game theory-based approach to determine the dynamics in the temporal evolution of Aβ aggregates along the pathways influenced by anionic fatty acid surfactants (*Ls*). Our rationale for such an approach is that the stochasticity and the often exclusive pathways of Aβ aggregation present a ‘win or loss’ scenario with respect to pathway adoption, governed by the concentration and phase-trasitions of the *Ls*. The mathematical analysis of this problem is taken up in two layers. The first is a six-species coarse-grained, reduced order model (ROM) while the second is a more detailed version of the model called, ensemble kinetic simulation (EKS) which captures the temporal kinetics of reactions at the atomistic scale. These models are partly validated by bulk kinetic and thermodynamic features using biophysical experiments. The simulations, supported by biophysical analyses, provide a temporal contour map along competing pathways, and present a unique perspective of otherwise unknown aggregate evolution along multiple pathways.

## 2 Materials and Methods

### 2.1 Reduced Order Kinetic Modeling (ROM)

The model presented here consists of a reduced order, comprising of only six species of Aβ that interact with the fatty acid surfactant, *L*, with the objective of understanding the complex dynamics of the system. Even with just six species, there are infinitely many rate regimes, most of which would be physically meaningless. Thus, only physically meaningful rate regimes derived from experiments and our previous studies (Ghag et al., 2013; Rana et al., 2017) were chosen and key parameters were vaired to understanding the dynamics.

A schematic of such a model is presented below. In this model, Aβ monomers can react with the pseudo-micelle *L* to create on or off-pathway oligomers. The system of chemical reactions in our model consist of the following,

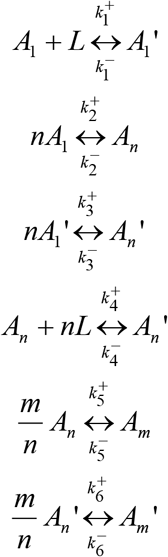

The non-prime species, *A*_1_, *A_n_*, and *A_m_* represent on-pathway Aβ monomers (*A_1_*) and oligomers (*A_n_ and A_m_*); whereas the prime species, 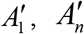, and 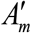, are the corresponding off-pathway species which are created through a reaction with the pseudo-micellesurfactant, *L*. The rate consants 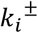(i=1-6) are indicated in the reaction schematic above where the ‘+’ represents a forward rate and ‘–’, a backward rate. These reactions were formulated based on experimental evidence demostrated earlier (Ghosh et al., 2016). In the computations to follow, for each species, *n* = 4 and *m* = 20 unless otherwise specified, which denotes the order of oligomer.

The reaction scheme is used to develop the corresponding kinetic model comprising of a system of six nonlinear differential equations. This system is then put into non-dimensional form. Using *A*_0_ as the characteristic concentration of monomers and 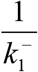 the characteristic time, we define the dimensionless species as follows

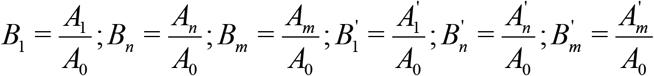

The reaction constants are similarly defined as follows:

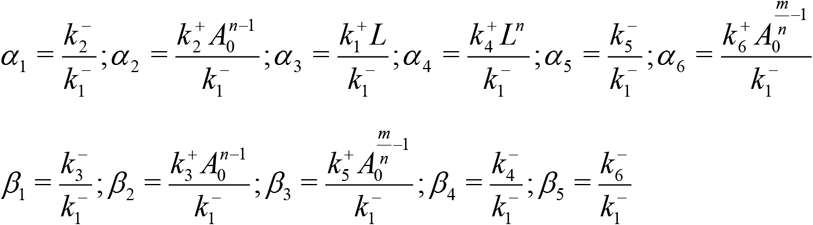

Note that both ∝_3_ and ∝_4_ have a factor *L* which is responsible for off-pathway aggregation. These two parameters serve as the *bridge variables* between on- and off-pathway species in the analysis which follows. Using the law of mass action kinetics, we formulate the dimensionless system of differential equations as follows

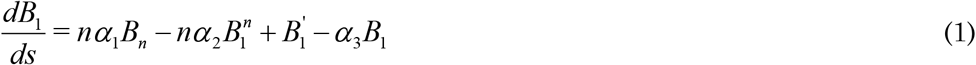

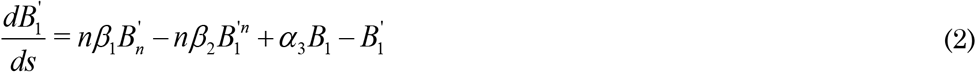

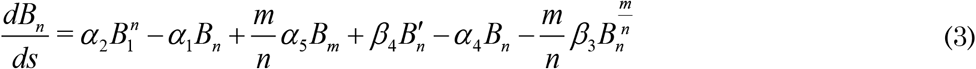

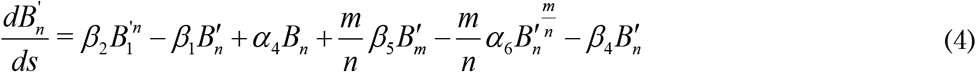

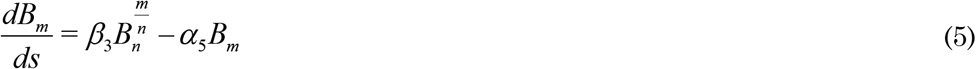

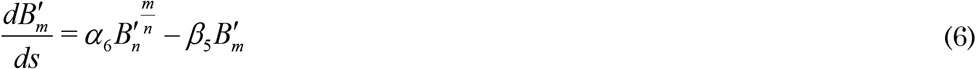

In this paper, we have primarily analyzed two models referred to as the *Base Model* and *Model 2* which are distinguished by the choice of fixed parameter values; i.e, the rate constant ratios in the purely on- and off-pathways. Our choices of the parameters in these two models are guided by prior work which have shown to capture the essential dynamics of the system, without loss of generality (Rana et al., 2017). In the base model, we set all forward rates (*α*_1_,*α*_2_,*α*_5_,*α*_6_) and bridge rates (*α*_3_,*α*_4_) equal to unity and all backward rates (*β*_1_,*β*_2_,*β*_3_,*β*_4_,*β*_5_) to 0.001 as previous mathematical models and experimental data have indicated (Ghag et al., 2013; Ghosh et al., 2016;). In the context of our model, a forward rate is defined as one that converts a smaller molecular structure into a larger structure, and backward being the reverse process. In model 2, we set all forward and backward rates to be equal to unity. *Ode 45* solver (Matlab) was used for our numerical computations.

**Figure1:**
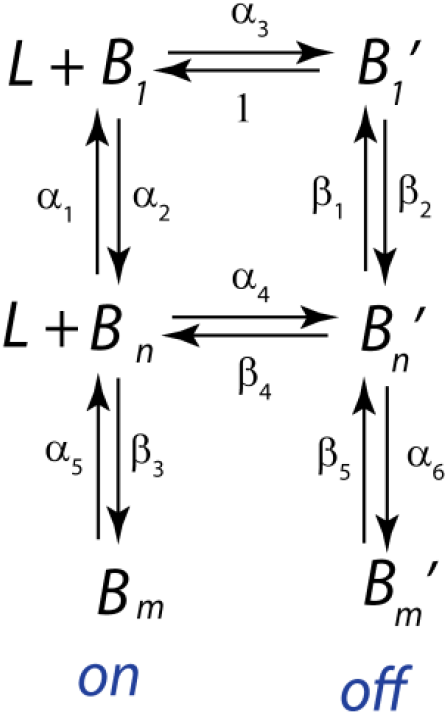
Schematic of on- and off-pathway aggregation model based upon the six-species reaction scheme described earlier.

### 2.2 Ensemble Kinetic Simulation (EKS)

To get more detailed insights into the switching behavior between on- and off-pathways, we formulate a combined off-on-pathway ensemble kinetic simulation (EKS) model. This EKS model has previously been applied for Aβ aggregation (Dean et al., 2018; Dean et al., 2017; Ghag et al., 2013; Ghosh, Datta and Rangachari, 2012; Ghosh et al., 2010; Lee et al., 2007; Rana et al., 2017) for both on-pathway and off-pathway. In this paper, we have extended our previous work by adding switching reactions considering off-to-on and on-to-off oligomer conversion. Please note, these switching reactions only take effect from a perturbation event such as changes in the concentrations of fatty acid, *L*. Specifically, dilution of *L* below its CMC triggers off-to-on switching while pseudo-micelleaddition triggers on-to off-switching respectively.

In this EKS model, we first considered a set of minimalistic reactions to represent the on-pathway, off-pathway, and its switching. Next, the flux for each reaction was computed. The system of differential equations of each species present in the reaction system were identified and solved using the ODE 23s solver (MATLAB). Below, we provide the reaction scheme while corresponding differential equations are presented in the Appendix.

I. *Reactions of on pathway: (considering A_12_ as F)*

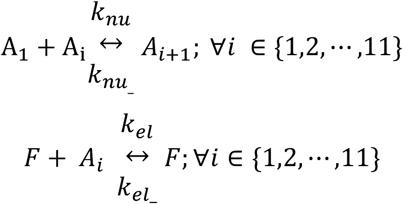
II. *Reactions of off pathway model:*

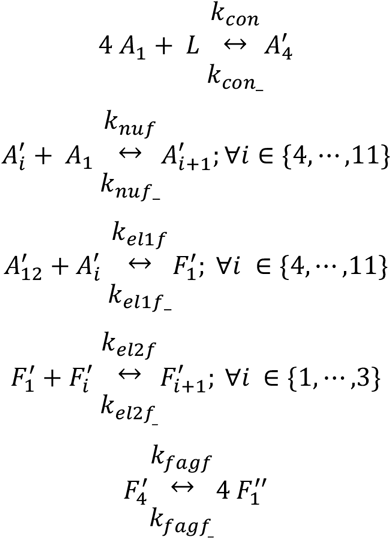
III. *On to off switching reaction:*

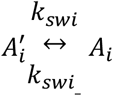

The corresponding flux for the reactions are given as:

I. *On pathway reactions flux*.

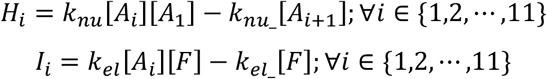
II. *Off-pathway reactions flux*.

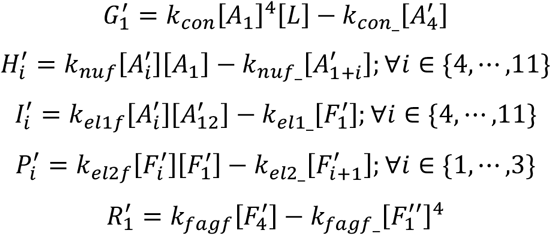
III. *Switching flux*.

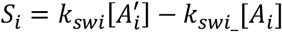

Here, and in the Appendix, *A_i_* denotes an on-pathway i-mer, *A’_i_* denotes an off-pathway i-mer, *L* denotes pseudo-micelles, *F* denotes post-nucleated on-pathway oligomers (termed as on-pathway fibrils; here *A_12_* is considered equivalent to *F* which corresponds to an on-pathway nucleus of 12mer that was previoulsy reported (Ghosh et al., 2016), *F’_i_* is an off-pathway oligomer, *signal* is the total ThT signal which is expressed as the sum of the on-pathway ThT signal (signal_on_) and the off-pathway ThT signal (signaloff) (as shown in Appendix; this uses an arbitrary mapping constant to map the total oligomer concentration to the experimentally observed ThT signal intensity). Note that in the EKS models, we consider the most general case where there can be switching between any on- or off-pathway oligomer of size *A_1_* to *A_11_*. Note that *A_12_* is considered as the nucleus, while all on-pathway oligomers beyond that are considered as on-pathway fibrils: *F* for the sake of simplicity. Similarly, smaller off-pathway oligomers from *A’_16_ – A’_23_* are considered as *F’_1_* and larger off-pathway oligomers are considered as *F’_i_*(*i= 1,…,4*) while a dissociation of *F’_4_* leads to the formation of *F’’_1_* which is a kinetically trapped off-pathway oligomer that does not aggregate further. The existence of such on- and off-pathway oligomers and the validity of our combined on- and off-pathway model (barring the switching reaction) has already been shown in earlier work (Powers and Powers, 2006; Rana et al., 2017).

### 2.3 Biophysical analysis

Synthetic wild-type Aβ42 was procured from both peptide 2.0 and Dr. Chaterjee’s laboratory at University of Mississippi were used in this study. Thioflavin-T (ThT), sodium dodecyl sulfate (SDS) and Lauric Acid (C 12:0) was purchased from Sigma-Aldrich (St. Louis, MO). Monoclonal Ab9 or Ab5 antibodies were obtained from University of Florida Center for Translational Research in Neurodegenerative diseases.

#### 2.3.1 Protein preparations

##### Preparation of Aβ42 monomers

Aβ42 peptide (1-1.5 mg) that was kept desiccated at −20 °C was dissolved in 50 mM NaOH and was allowed to stand at room temperature for up to 45 minutes. The dissolved peptide was then fractionated on a Superdex-75 HR 10/30 size exclusion chromatography (SEC) column (GE Life Sciences) on a BIORAD FPLC system that was pre-equilibrated with 20mM Tris at pH 8.00, to separate any preformed aggregates as previously reported ^1^. Fractions were collected at a flow rate of 0.5 mL/min and stored at 4 °C until use. The concentration of the monomeric fractions was calculated using a Cary 50 UV-vis spectrophotometer (Varian Inc.). The molar extinction coefficient of 1450 cm^-1^ M^-1^ at 276 nm was used (www.expasy.org). Monomers were used within one day after fractionation.

##### On- and off-pathway aggregation reactions

The on-pathway aggregates were initiated using 40 μM monomeric Aβ42 in 20 mM TrisHCl, 50 mM NaCl at pH 8.0 incubated under quiescent conditions at 37 °C with and 0.01% NaN_3_. Off-pathway reactions were initiated using 25 μM monomeric Aβ42 in the same buffer incubated with 50mM NaCl and 5mM sodium laurate (C12 FA) in 20mM Tris, pH 8.00, as reported previously (Powers and Powers, 2008; Rana et al., 2017).

#### 2.3.1 ThT Fluorescence aggregation assay

For on-to-off pathway switching reactions, to 150 μL, 50 μM Aβ reactions incubated in buffered alone, a 50 μM thioflavin-T (ThT) solution in the same buffer was added and fluorescence emission (λ = 482 nm) was collected using microplate reader (Biotek Synergy Microplate reader) at 37 °C using an excitation at 452 nm. A 5 mM sodium laurate (C12 fatty acid) sample pre-equilibrated with 50 μM ThT was added to the reactions at 3, 8, and 24 h to initiate switching of pathways. The data was collected at ten-minute time intervals. For off-to-on-pathway switching reactions, the 150 μL, 50 μM Aβ reactions pre-incubated in the presence of 5 mM sodium laurate were diluted 5- or 10-folds at 5 and 10 h using buffered 50 μM Aβ monomers and 50 μM ThT such that only the fatty acid concentration is dropped below its critical micelle concentrations. Appropriate blank reactions were monitored simultaneously and were corrected before data processing.

#### 2.3.2 SDS-PAGE and immnunoblotting

Aliquots of the reactions were mixed with sample buffer comprising 1% SDS (1X Laemmli sample buffer) and loaded on a precast 4-12% Biorad gel. For calibration, pre-stained molecular weight markers (Invitrogen inc) were used. The gels were then electroblotted on 0.45 um nitrocellulose membrane (GE Life Sciences). The blots were then heated in microwave for 1 min and were blocked with 5% non-fat dry milk solution with 1% Tween-20 in PBS for 1.5h. Subsequently, the blots were probed with monoclonal antibody Ab5 or Ab9 (1:1000-1:2500 dilutions) which bind to residue 1-16 of Aβ. Anti-mouse horseradish peroxide was added to the blot and the blot was developed with ECL reagent (Thermo Fisher Scientific) and imaged with a Bio-Rad Gel Doc system.

## 3 Results

### 3.1 ROM indicates switching behavior between pathways is dictated by the dynamics of equlibrium stability and bridging

#### 3.1.1 Equilibrium Points

In order to study the stability of the system we rely on experimental data to determine rate constants which in turn determine our equilibrium, or fixed point, concentration levels (Ghag et al., 2013). We are especially interested in the non-zero equilibrium states of each species, i.e. concentrations that would simultaneously make equations (1)–(6) disappear for a given set of parameter values. Numerical computations indicate that concentration of species continue to fluctuate over time for our models, but in all cases these fluctuations were within 0.1% of previous levels for *t* greater than some critical time, which we considered acceptable as equilibrium. The equilibrium values, were also confirmed through Matlab’s *fsolve* function. In all ROM computations discussed in this paper, the initial conditions were taken to be *B*_1_(0) = 1 with all other species set initially at zero. As seen from the figure 2a and 2b, as time increases, the concentration levels exhibit asymptotic behavior and each species eventually achieves equilibrium. The time to reach equilibrium is sensitive to the choice of rate constants; the base model takes much longer to reach steady state than the model 2. Also, in all cases analyzed the fibril concentrations *B_m_* and 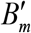 take the longest to reach equilibrium.

**Figure 2:**
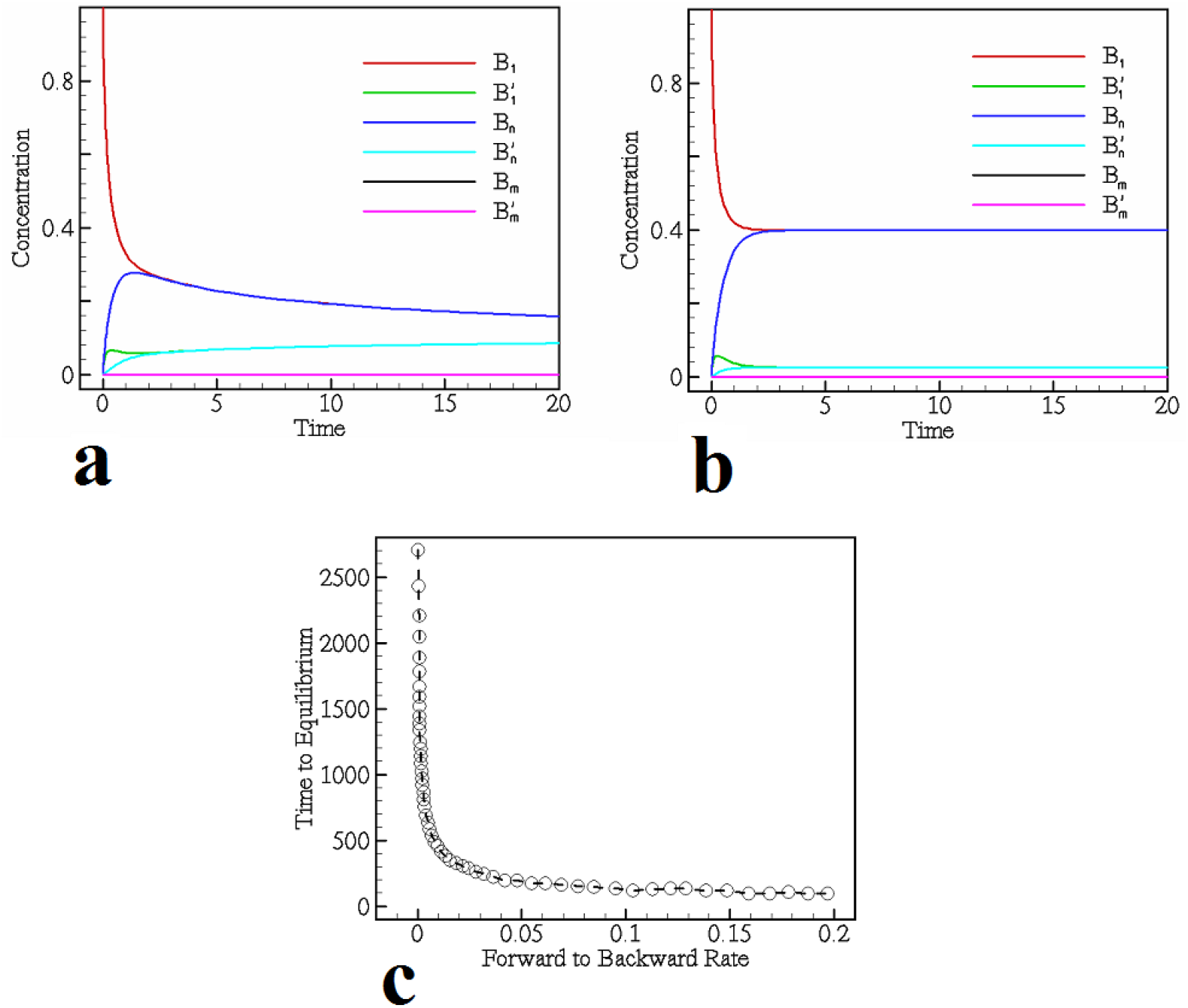
Panels (a) and (b) show sample solutions of the *base model* and *model 2* corresponding to equations (1)–(6) for the ROM. The different colors in both panels correspond to the evolution of the six different species, indicated in the figure legend. Panel (c) shows the equilibrium time as a function of the ratio of backward to forward rates for the *base model*. Clearly, as the ratio of forward to backward rate constants increase, as in pathalogical cases, the time to equilibrium declines.

Due to low forward rates, the concentration size of *B*_1_ stays high and stable throughout, but there are periodic, large percentage changes in the concentration of 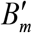. We analyzed the concentration patterns of both *B*_1_ and 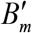 over this period and their growth and declining patterns are a mirror image of each other: periods of increase in *B*_1_ were accompanied by declines in 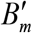. From our equilibrium analysis of these models we found that when the ratio of backward to forward rates is close to 1, the model settles at equilibrium more quickly than when there are large differences in the magnitude of backward and forward rates (see figure 2c). A power-law regression indicates that time to equilibrium (indicated *t_eq_*) varies with the ratio of forward to backward rates (*r*) according to *t_eq_* ≈ *r*^−0.615^.

The impact of varying *n* and *m* in both models was also investigated. In these cases, increasing *n* and *m* had the effect of increasing *t_eq_*. It appears that the larger the oligomer size, the higher the power of the non-linear terms in the governing equations, the greater the potential for over- and under-shoot as the model evolves over time, thus, taking longer to achieve equilibrium. However, model 2 does not reveal such an increase in *t_eq_* as noted earlier.

#### 3.1.2 The Bridge Parameters

The key parameters in our model are ∝_3_ and ∝_4_ which serve as "bridge parameters" that govern the reaction dynamics between on- and off-pathway. We verify the effect of varying both on species formation while holding all other reaction rates constant (figure 3). When increasing both ∝_3_ and ∝_4_, we see a direct increase in the ratio of off-pathway species to on-pathway species. Because 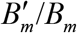 is not directly governed by the bridge variables, it is slower to react to changes along the bridges, but eventually exhibits what appears to be exponential or power growth at higher values of the bridge variables (see figure 3a). This is most likely due to the fact that 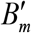 formation is dependent upon and ∝_3_ and ∝_4_, so that increasing ∝_3_ and ∝_4_ eventually significantly impacts 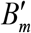.

**Figure 3:**
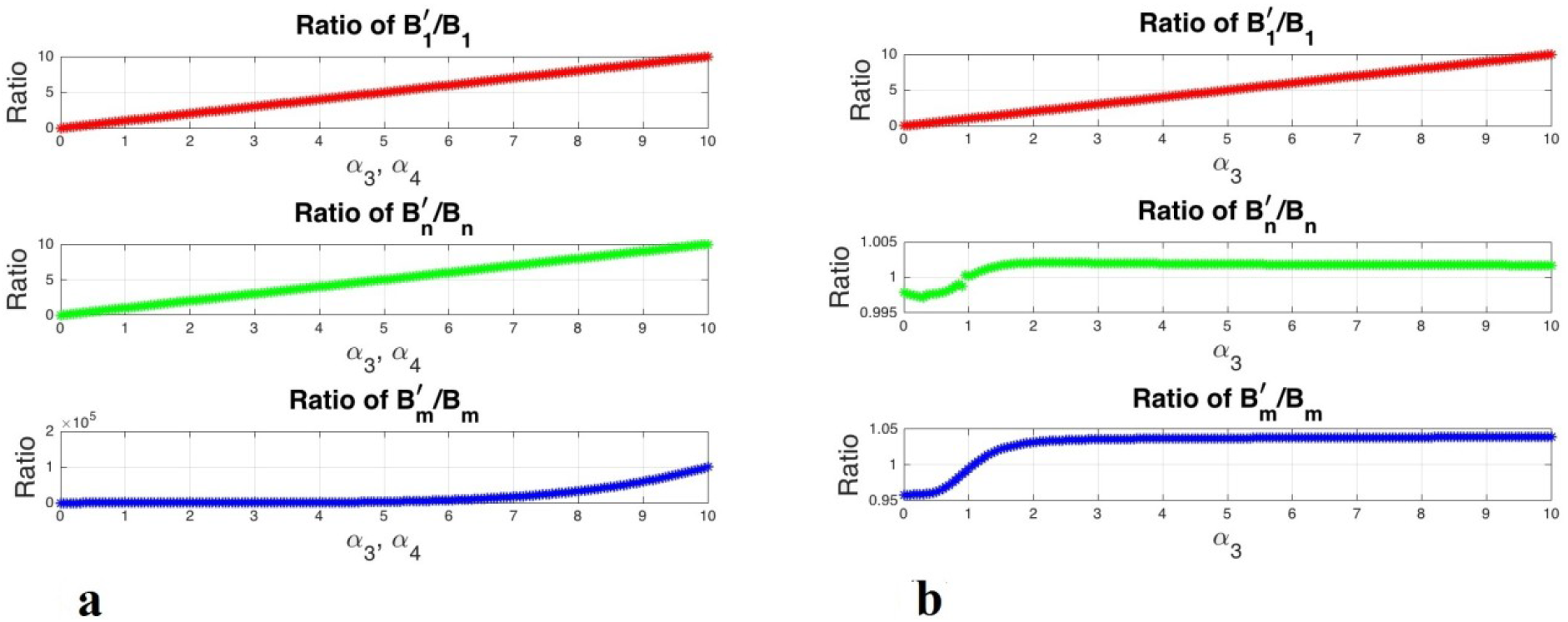
Concentration ratios of like species between the two pathways as a function of ∝_3_ and ∝_4_ for the base model. The panel (a) shows these ratios as a function of both the bridge parameters while in panel (b) ∝_3_ is varied while holding ∝_4_ fixed. This figure shows the impact of the bridge parameters upon specific oligomers in the reaction pathways, indicating that the bridges between larger oligomers play a more significant role in the pathway dynamics and competition.

Figures 3a and 3b underscore the importance of the bridge variables. Interestingly, if we leave ∝_4_ unchanged and simply increase∝_3_, there is limited flow-through from 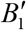 to 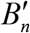 and 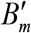: their ratios to the non-prime species increase slightly above unity, but cease to grow from there even as ∝_3_ continues to increase. Therefore, the bridge reaction 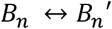 is critical in the formation of the larger oligomers, i.e. the *n* and *m* species. The ROM modeling therefore reveals that bridges between larger oligomer are more significant than the ones across monomers in terms of promoting off-pathway fibril formation.

Additional tests were performed to verify conditions for any species to outperform others by appropriate choice of the rate constants. Forcing *B*_1_ to outperform, for instance, is just a matter of greatly reducing or shutting off all the forward reactions. For species further down the reaction-network, we are required to increase forward reactions to get the desired out-performance. In the case of 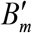, we were able to get out-performance of this species in absolute terms by dramatically increasing the forward reaction rates ∝_3_, *β*_2_, and ∝_6_ by an order 10^4^. Out-performance by *B_m_* could also be achieved in a similar manner. Such an exercise can be be significant in helping identify pathologies and shows the robustness of the network under standard reaction rates.

#### 3.1.3 Linear Stability Analysis

A linear stability analysis was conducted to confirm the conditions under which equilibria are stable and the sensitivity of these solutions to the parameters in this problem. We use the variables *X*_1_, *Y*_1_, *X_n_, Y_n_, X_m_* and *Y_m_* to represent the concentrations of the various perturbed species while the equilibria for the monomers and oligomers from the two pathways are indicated by means of an ‘e’ in the subscript (i.e. *B_k,e_* represents the equilibrium concentration for the oligomer of size *k*). The central idea behind the stability analysis being that a stable equilibrium requires that the perturbed quantities eventually vanish, under certain conditions.

The linearized system of equations for the perturbations is given by

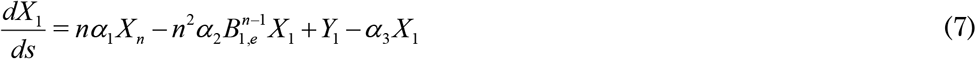

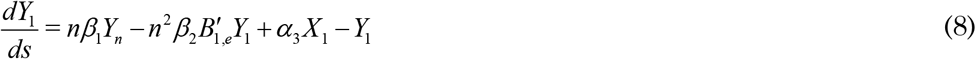

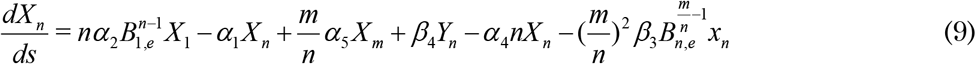

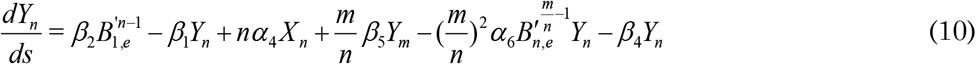

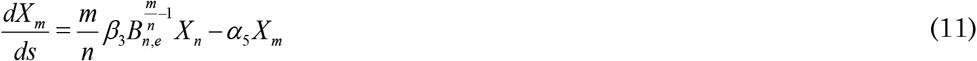

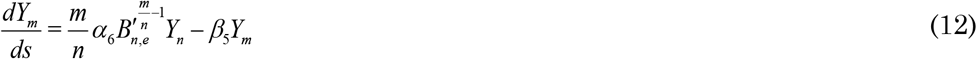

The equations can be represented in operator form and solved for its eigenvalues (denoted *λ_i_*, where *i*=1-6). Of key interest is the effect of the bridge parameters, ∝_3_ and ∝_4_, and their effect on the stability of each model. A sampling of this effect is captured in figure 4 which depicts the contour plots of the eigenvalues of the base model for 0 ≤ *α*_3_,*α*_4_ ≤ 2 for the special case when *n* = 8 and *m* = 4 0. In this figure, the lighter shades depict regions of low stability while the darker ones are more stable. The eigenvalue *λ_i_* corresponds to neutral stability. Overall, we find that the stability profile for equilibria corresponding to the base model does not change much for variations in values of *n* and *m*. The stability picture for the base model, however, is significantly different from that of model 2. In the base model, one of *λ*_4_ or *λ*_5_, is always zero for all values of ∝_3_ and ∝_4_, while for model 2, we observe switching behavior between *λ*_2_ and *λ*_4_, i.e. the model 2 shows greater sensitivity to the values of the bridge parameters.

**Figure 4:**
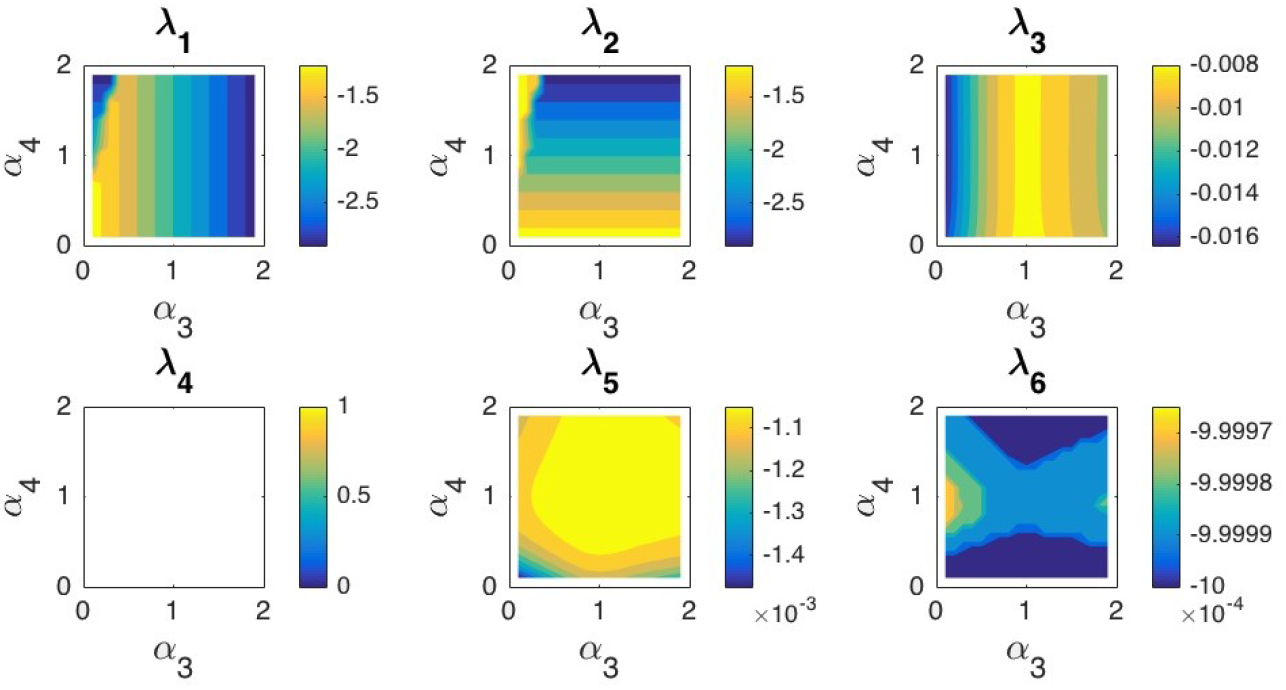
*λ_i_* as a function of ∝_3_ and ∝_4_ for the case of *n* = 8, *m* = 40. The deeper blue shade represents the more stable regions. The eigenvalue *λ*_4_ is zero for all values of the parameters indicating neutral stability.

**Figure 5:**
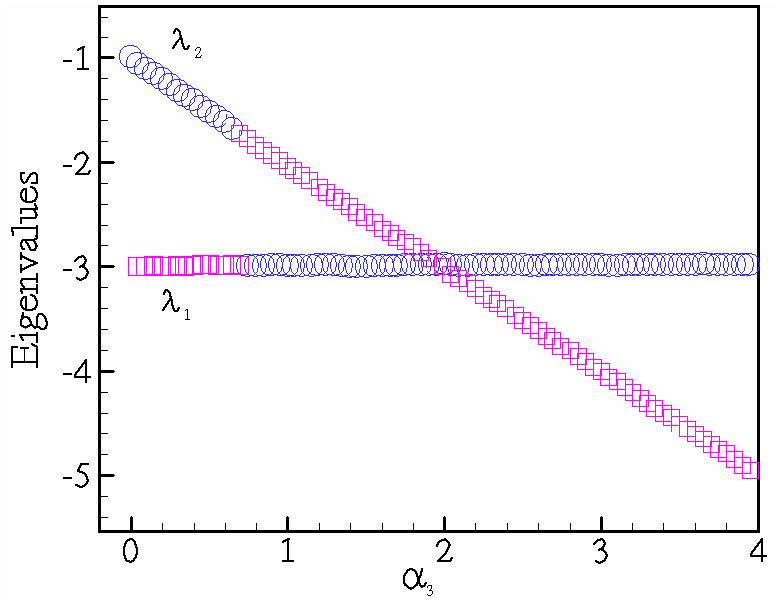
A sample case of eigenvalues depicting switching and crossing as a function of the bridge parameter ∝_3_. The eigenvalue *λ*_2_ is denoted in blue while eigenvalue *λ*_1_ is indicated in lavender.

We distinguish two different characteristic effects, namely *switching* and *crossing* of the eigenvalues as the two bridge parameters are varied. The switching indicates a sudden, drastic change in behavior of the species, where the course of domination of one species over other is abrutly reversed while the crossing is a more gradual version of this shift. In previous work (Rana et al., 2017), the switching has been compared with a sort of *transcritical-like bifurcation* in the system. The Table 1 shows the switching and crossing points for eigenvalues *λ*_1_ and *λ*_2_ as ∝_3_ and ∝_4_ vary. As can be seen, there is crossing where ∝_3_=∝_4_, whereas switching has an exponential relationship between the two parameters. A regression model indicates that switching occurs according to ∝_4_ ≈ 2.04 × 1.55^∝_3_^ with *R*^2^ = 0.953.

**Table 1.**
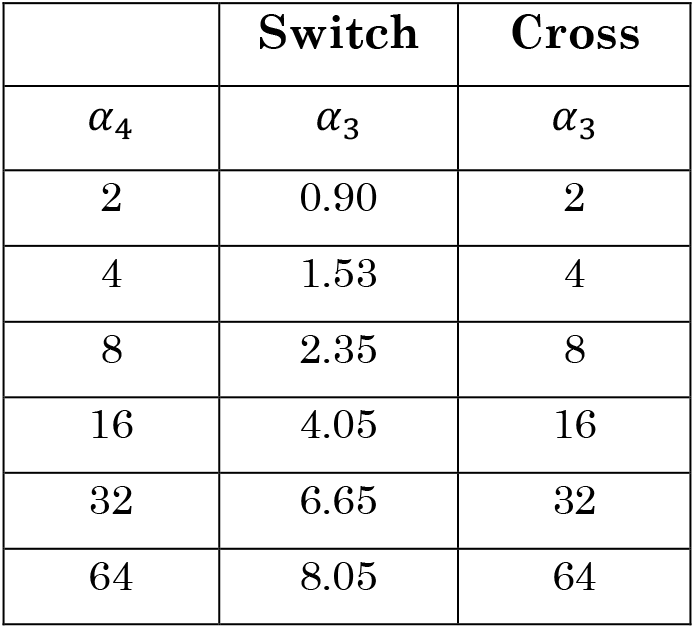
The table shows switch and cross points *α*_3_ of as a function of *α*_4_.

Simulations for other reaction rate regimes over a larger range of values for ∝_3_ and ∝_4_(greater than 2) showed the switching and crossing to persist, as indicated by Table 1. In previous studies with lower-dimensional models (Rana et al., 2017), we have seen such switching to occur as well which appears to be indicative of the sensitivity of the system to the various pathways and species in the model and activation of one of these pathways under appropriate conditions. For the base model, with *n=2* and *m=4*, we have *λ*_3_ =0 for low ∝_3_ and ∝_4_, but as we increase these two parameters, *λ*_2_ vanishes and then finally, with further increase, *λ*_4_ goes to zero. Thus, for any rate environment, the stability of the system near the point of equilibrium was found to be neutrally stable for suffiicently large values of the bridge parameters.

The impact of species-size upon stability was also examined by studying the cases of (*n,m*) equal to (2,4) and (8,40) in addition to the standard case (4,20). Overall, in the case of the base models, no switching of eigenvalues was observed for a given species size environment as ∝_3_ and ∝_4_ were varied. For instance, in the base model, *λ*_5_ is zero for all values of and when *n* = 2 and *m* = 4, and is zero, when *n* = 8 and *m* = 40. However, for model 2, there is switching between *λ*_2_, *λ*_3_, and *λ*_4_ when *n* = 2 and *m* = 4, whereas when *n* = 8 and *m* = 40, we observe switching between *λ*_1_, *λ*_2_ and *λ*_3_. In general, as *n* and *m* are increased, the overall magnitude of stability increases, i.e. *the larger the species, the more stable the individual oligomer, and also the overall system, appears to be*.

#### 3.1.4 Game theoretic approach to understanding pathways

A significant and novel approach to the problem lies in analyzing the model equations (1)–(6) from a game-theoretic point of view. The figure 6d provides a schematic of the four equilibrium pathways that our model can achieve, each sensitive to the choices of parameters ∝_3_ and ∝_4_. Consider the Figure 6d, from top-left, clockwise. The first schematic highlighted in red is strictly on-pathway, where the non-prime species “win”. The next highlighted in blue is strictly off-pathway, with all off-pathway species winning. The paths indicaated in yellow and green are a mixture of on/off-pathway. Our computations were conducted by varying ∝_3_ and ∝_4_ between 0 and 2 in increments of .02, resulting in 10,000 discrete points. The figures 6a and 6b show a phase-diagram for the base model and model 2, respectively.

For the base model, (*α*_3_,*α*_4_) = (1,1) is a critical point at which the concentration of on- and off-pathway species are equal. As we vary ∝_3_ and ∝_4_, from this point we begin to see dominance of one set of species or pathways over another. Notable too is the fact that the boundaries between the different equilibrium states are almost linear: the line *α*_4_ =1 determines the switching between on- and off-pathway dominance of *n* and *m* species, and the line *α*_3_ =1 determines the switching between on- and off-pathway dominance of monomers. The table in figure 6c shows the equilibrium states as a function of ∝_3_ and ∝_4_ in the form of a payoff matrix. The *Nash Equilibrium*, lies where on-pathway species concentrations are equal to off-pathway species concentrations.

A similar computation was performed for model 2, which is shown in the figure 6b. Once again, the point (*α*_3_,*α*_4_) = (1,1) is a critical point, however, unlike in the base model, it doesn’t strictly define outperformance of *B*_1_ over 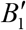 and vice versa; we still see *B*_1_ outperform 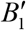 for *α*_3_ > 1 and low *α*_4_, and 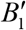 outperform *B*_1_ for *α*_3_ < 1 and high ∝_4_. The major difference is that the red and blue regions representing the strictly on-pathway and strictly off-pathway respectively, increase at the expense of green and yellow (mixed pathways).

More importantly, the range of points over which on-pathway wins, gets bigger when backward rates are on par with forward rates. For *α*_3_ <1, by increasing *β*_1_ and *β*_5_ we are effectively shutting off the 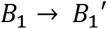 bridge towards off-pathway aggregation, hence the increase in red in the upper left of the figure 6b. The reverse happens in the lower right for *α*_4_ < 1 as we get greater off-pathway aggregation up to a point. Despite increasing *α*_3_ above 2, however, if we reduce ∝_4_ we continue to see a thin band of dominance of *B_n_* and *B_m_* over their respective primed, off-pathway species. Once again, this shows that the 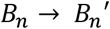 bridge is more critical for off-pathway aggregation of *n* and *m* species than the monomer-bridge. If we reduce *α*_4_ we still get on-pathway dominance of *n* and *m* species even for high ∝_3_. Thus, it is difficult to control the off-pathway aggregation of *n* and *m* species by tweeking the 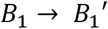 bridge.

**Figure 6:**
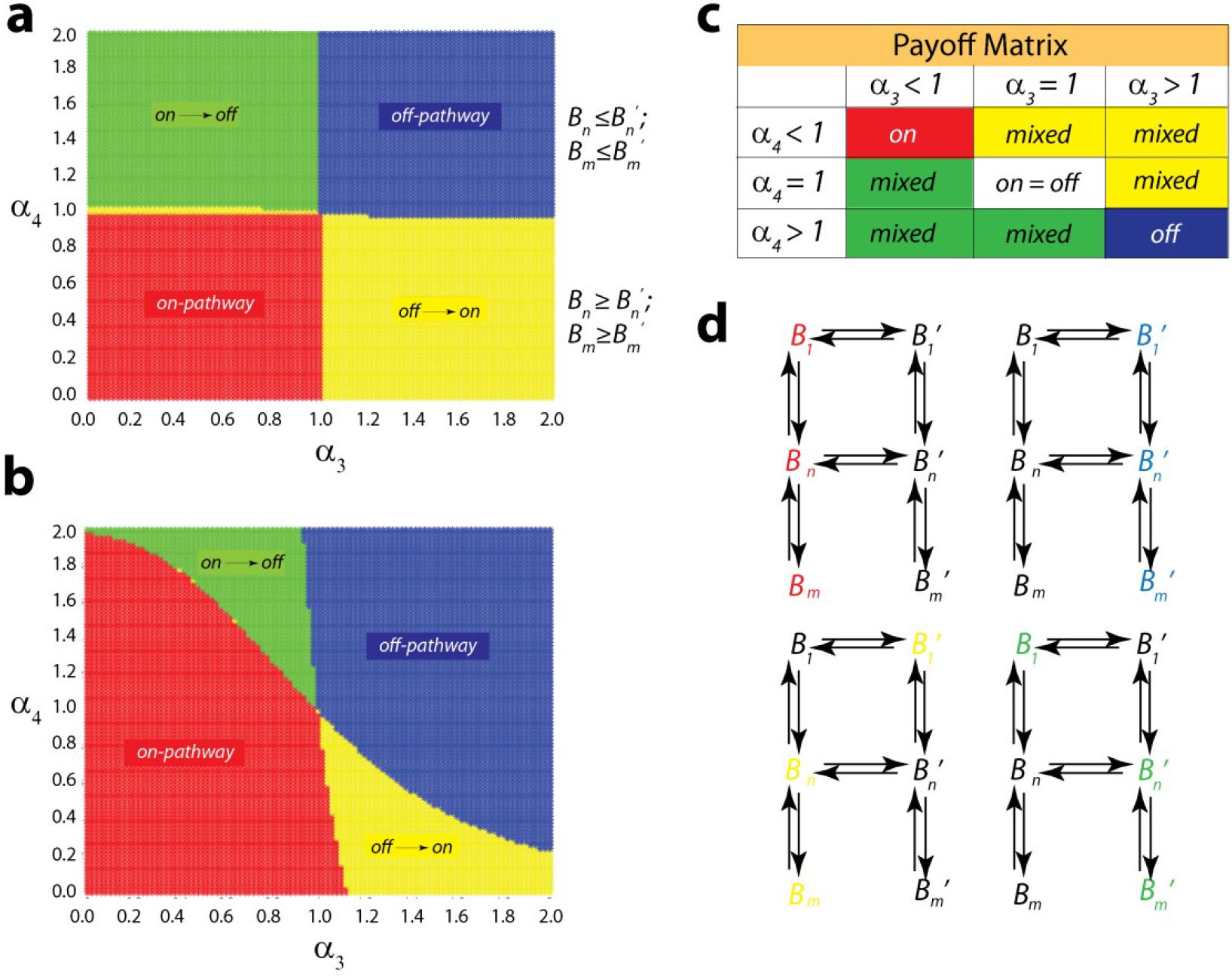
Aggregation pathways as a function of *α*_3_ and *α*_4_. The panels (a) and (b) depict a contour plot of the dominant species as a function of the bridge parameters for the base model and model 2, respectively. Panel (c) shows the pay-off matrix for figure (a) depcting the various conditions for domination. Panel (d) depicts the pathway diagram indicating the dominant sub-path for specific choices of bridge parameters, correspoding to figures (a) and (b).

### 3.2. Biophysical evidence for the switching of aggregation pathways

The effect of fatty acids on Aβ42 aggregation has been well established in Rangachari laboratory (Dean et al., 2018; Dean eat al., 2017; Rangachari et al., 2007). Specifically using sodium laurate at concentrations near and well above its CMC, the surfactant was able to modulate Aβ42 aggregation toward off-fibril formation pathway that was populated by low molecular weight oligomers. At concentrations well below CMC, the fatty acid adopted a on-fibril formation pathway (Kumar et al., 2011). To experimentally assess the swithchig of pathways from on-to off-pathay and vice versa by modulating *L* concentrations, kinetic rate differences in aggregation was investigated using ThT dye.

**Figure 7:**
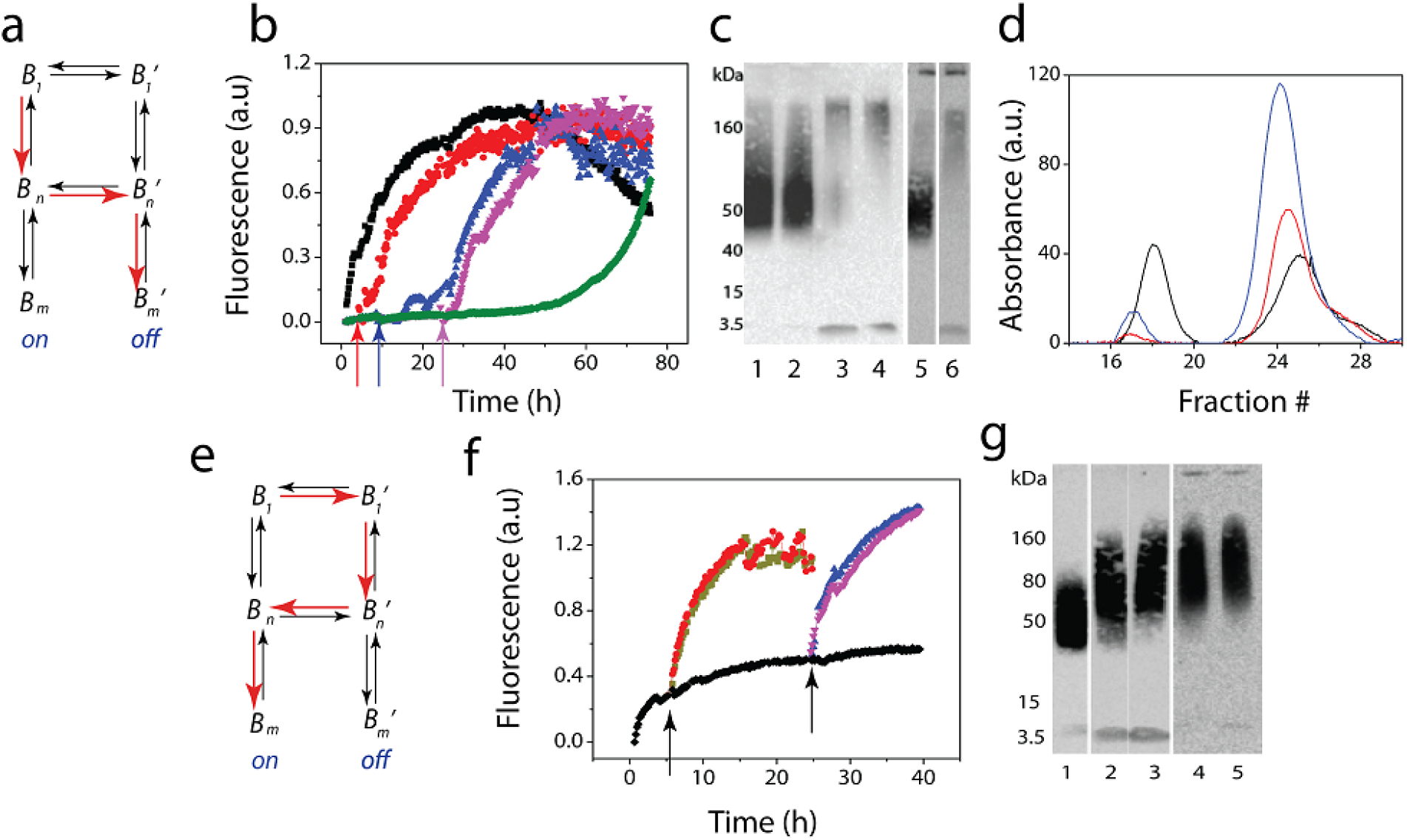
Experimental verification of switching of pathways. **(a)** schematic representation of on-pathway switching (red arrows) of on-(*B_n_*) to off-pathway (*B_n_’ and B_m_*) on addition of C12 FA **(b)** ThT kinetics of the on-to-off transitions probed by the introduction of fatty acid at 3 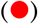, 8 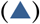 and 24 h 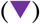 time points, along with the controls with no fatty acid 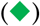 and with C12 FA introduced at 0 h 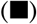. **(c)** Immunoblots for the corresponding reactions: addition of 5 mM at 3h (lane 1); addition of 5mM at 8h (lane 2); 3h, 8h and 24 h buffer controls (lanes 3, 4 & 5), and addition of 5mM at 24h (lane 6). **(d)** SEC fractionation of the reaction before the addition of fatty acid at 24h (blue), involving the addition of 5 mM C12 fatty acid (black) at 24 h to the sample and control without fatty acid (red), after subsequent incubation for 24 h at 37 °C. **(e)** schematic representation switching of off- (*B_n_*’) to on-pathway (*B_n_’ to B_n_*) on dilution of the fatty acid below its critical micelle concentration (f) ThT kinetics monitored by the removal of 5 mM fatty acid on the sample incubated with Aβ by diluting with buffer either 5-(red or purple) or 10-folds (green or blue) at 5 h 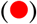 and 24 h 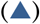, respectively. The control without dilutions is shown in black; 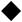. (g) Immunoblot of off-pathway oligomer control generated in the presence of 5 mM fatty acid at 24h (lane 1); 5- and 10-fold dilutions at 5h respectively (lanes 2 & 3), and 5- and 10-fold dilutions at 24h respectively (lanes 4 & 5).

Switching of on-to off-pathway (depicted schematically in Figure 7a) was initiated by the addition of 5 mM C12 FA to 25 μM Aβ42 buffered in 20 mM Tris, 50 mM NaCl at pH 8.0. Addition of C12 FA resulted in an increase in ThT fluoresnce without ann observable lag time (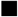; Figure 7b). In contrast, Aβ42 in the absence of C12 FA showed a lag phase of ~ 50 h before an increase in ThT fluorescence was observed (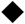; Figure 7b). This behavior in the presence of C12 FA has been as previously observed to generate 12-24mer oligomers of Aβ along off-fibril formation pathway (Dean eat al., 2016). In order to evaluate the propensity of bridging from on-to off-pathway, 5 mM C12 FA was added to the control Aβ42 reaction after 0h (positive control 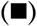, 3 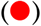, 8 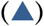 and 24 h 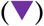. Each of such incubation resulted in an exponential increase in ThT fluorescence suggesting switching of pathways from on-to off (Figure 7b). Analysis of these samples was also perfomred using a partially denaturing gel electrophoresis (low SDS and no boiling) and immunoblotting (Figure 7b). Injections of C12 FA at 3 and 8 h show the presence of 48-60 kDa band corresponding to 12mer oligomers (lanes 1 and 2, respectively) as compared to the corresponding controls generated upon adding buffer in place of C12 FA (lanes 3 and 4), which show monomers and some on-pathway aggregates. This suggests that off-pathway oligomers are generated (Figure 7c). Similarly FA injected after 24 h and its corresponding control is shown in lanes 5 and 6, respectively, which shows evan after 24 h, C12 FA is able to induce the formation of oligomers to certain extent, with clearly observable emergence of some on-pwathay fibrils. These results parallel those observed by ThT fluorescence (Figure 7b).

To further quantify the extent of bridging, the aggregates generated after the 24 h injection of C12 FA (or buffer for the control) were fractionated by size exclusion chromatography (SEC), after an additional 24 h incubation (Figure 7d). Prior to fractionation, the samples were centrifuged at 18×000 g for 20 min to remove any high molecular weight fibrils, and the supernatant was loaded on to the column. After 24h, the control in the absence of C12 FA show a small peak near the void volume at fraction 17 and a monomer peak at fraction 24 (blue; Figure 7d). Fractionation of the control reaction at 48h (after injection of buffer at 24h) showed a diminished peak at fraction 17 and a reduced monomer peak at fractionn 24 (ref; Figure 7d). The reduction in the monomer peak correlates to they being consumed during aggregation. Similar reduction in the aggregate peak between 24 and 48 h can be explained by the fact that many have formed fibrils that are centrifuged out. On the other hand, fractionation of the sample after 48 h with the injection of 5 mM C12 FA at 24h, showed larger peak at fraction 18 and a reduced monomer peak at fraction 25 (black; Figure 7d). This suggests two possibilities; a) the unreacted monomers adopt off-pathway upon introduction of C12 FA, and/or b) the pre-formed aggregates along the on-pathway are re-routed back through the off-pathway, in other words, switching. A more detailed analysis on this is discussed later in the manuscript.

To assess a similar switching of pathways from off-to on-pathway (schematically depicted in Figure 7e) was performed again using the established C12 FA kinetics. Incubation of 5 mM C12 FA shows an exponential increase in ThT fluorescence (black 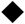; Figure 7f). To effect switching of off-to on-pathway afte certain time periods, the sample was diluted 5- and 10-folds such that the effective concentration of C12 FA drops to 1 and 0.1 mM, which are well below the CMC of the surfactant. It is well established that well below CMC, Aβ aggregation is augmented (Kumar et al., 2011), and therefore, dilutions of 5 mM C12FA must induce a faster rates of aggregation. When dilutions were introduced, at 5 and 24h time points (arrows; Figure 7f), appropriately blank subtracted data showed an increase in ThT fluorescnce as expected for both dilutions suggesting the switching of off-to on-pathway (Figure 7f). Partially denaturing gel electrophoresis and immunoblotting further confirmed the switching. The five- and ten-fold dilitions resulted in an increase in the molecular weight of the aggregates including the formation of fibrils both at 5 and 24 h, respectively (lanes 2-5; Figure 7f) as compared to the sample in 5 mM C12 FA (lane 1).

### 3.3 EKS models validate the game theoritic approach in elucidating the dynamics of a competing aggregation pathways

#### 3.3.1 Parameter estimation

In the EKS model, we have four on-pathway rate constants (namely, *k_nu_, k_nu__, k_el_, k_el__*), 10 off-pathway rate constants (namely, *k_con_, k_con__, k_nuf_, k_nuf__, k_el1f_, k_el1f__, k_el2f_, k_el2f__, k_fagf_, k_fagf__*), and two off-on switching rate constants (note that the forward and backward rate constants of switching each oligomer was considered the same leading to only two switching parameters that need to be estimated, i.e., *k_swi_, k_swi__*). Additionally, we also need to estimate two additional constants: *p* (which is simply a mapping constant that distinguishes the contributions of on-pathway oligomers from off-pathway oligomers to the ThT signal) and pseudo-micelle concentration (concentration of the fatty acid near its CMC denoted by *L*). Following our published model in (Rana et al., 2017), the pseudo-micelle concentration was additionally estimated and not calculated directly from the FA concentration values at the CMC since precise concentrations of pseudo-micelle are only a fraction of total fatty acid concentration that are difficult to determine experimentally. This increased the number of parameters that need to be estimated to 18 from the EKS simulations. Fortunately, our on-pathway and off-pathway rate constants can be estimated separately using the respective control data. This makes it easier to estimate the remaining four rate constants (i.e., the two off-on switching rate constants *k_swi_, k_swi__*, the mapping constant *p* and the pseudo-micelle concentration *L*) from this off-on switching dataset by significantly reducing the number of free parameters. A large parameter space from 10^−6^ to 10^8^ units with multiples of 10, was swept, to estimate the value of each of the two switching rate constants. Similarly, the pseudo-micelle concentration was varied from 0.01 to 1 unit, and *p* was varied from 10^5^ to 10^8^ units. The estimated parameter values corresponding to the best fits are shown in the Table A1 in the Appendix. The benchmarked on- and off-pathway rate constants (estimated separately from control data), were used to estimate the switching rate constants and obtain a global fit to the experimental ThT curves and monomer ratio values estimated from SEC measurements. The average R^2^ of the off-to-on data is 0.974 and that of on-to-off data is 0.981.

#### 3.3.2 Numerical Results

The switching rate constants were very sensitive specifically in the off-on dataset. The experimental data could not be fit in the absence of the switching rate constants and only a handful of switching rate constant combinations allowed an acceptable fit; the switching rate constants corresponding to the best fit to the experimental data are reported in the Table in the Appendix. This directly proves the switching of off-pathway oligomers to on-pathway oligomers through the switching pathways due to the dilution of the system. The EKS simulations were conducted in the same way as the experimental set-up. For the off-to-on switching (Fig 8), first, combined off- and on-pathway simulations were executed, upto the switching time-point (of 5 or 24 h); all oligomer concentration were noted until this point and they were then recalculated based on the amount of dilution at the switching time-point from the experiments. These altered concentration levels for each oligomer were next considered as the initial concentration of the combined off- and -on pathway simulation. Note that the second phase of the off-to-on dataset (Fig 8a) does not show any lag time as can be seen in a usual unseeded on-pathway system. Our model predicts a large conversion of off-pathway species to on-pathway oligomers which results in a rapid formation of on-pathway fibrils (denoted by *F*).

For the on-off dataset (Fig 8b), stand-alone on-pathway simulations were executed exclusively up to the switching time-point (24-h) and the current oligomer concentration were noted. These concentrations were then used to restart the combined on- and off-pathway simulation in addition to the pseudo-micelle concentration (that was also estimated in the parameter search step as an independent variable). Surprisingly, we found that the on-to-off pathway dataset could be fit to our model both considering the switching rate constants, and also in the absence of switching rate constants generating comparable *R^2^* values; in other words, the switching rate constants had low sensitivity to the on-off experimental dataset. Probably as the on-pathway reactions are slow, very little on-pathway oligomers are formed at the switching time-point; as a consequence, this made the switching reaction flux slower than the previous case of off-on switching system resulting in overall lower sensitivity of the switching rate constants to the ThT data points from the experiments. While this precludes precise characterization of on-off switching, we do observe an overall decrease in fibril concentration compared to control data showing at least a qualitative impact of the switching reactions that convert the on-pathway oligomers into off-pathway species.

**Figure 8:**
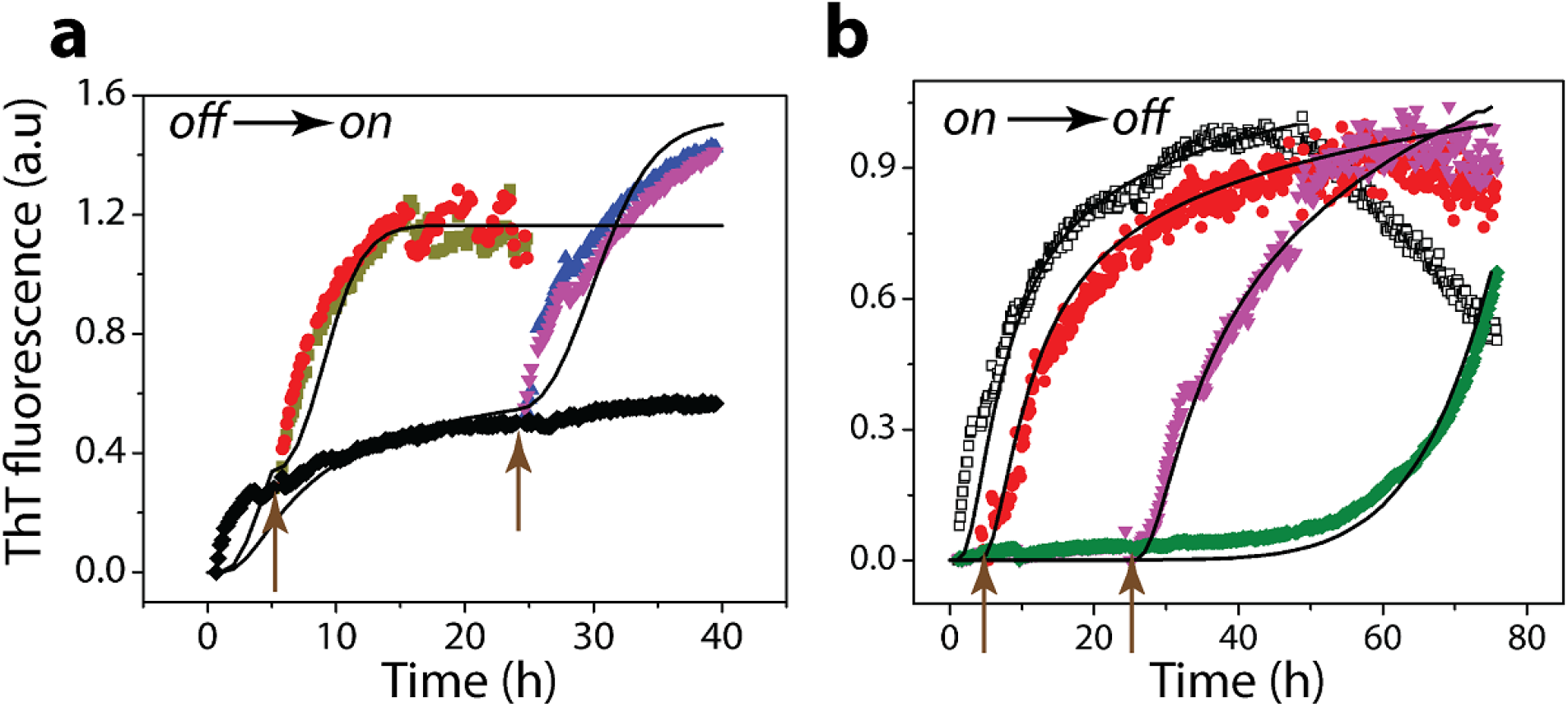
Correspondence between experimental results and EKS models on switching of pathways. a and b) Experimental data (scatter data) on on-to-off and off-to-on pathways reproduced from Figure 7b and 7f, respectively. Models based on EKS are shown as black lines. The intervention time points of 3 and 24 h (for (a)), and 5 and 24h (for (b)) are shown as arrows.

## 4. Discussion

The data presented here is a first attempt in deciphering the complex phenomenon of protein aggregation pathways using a classical game theroetic, Nash equilibrium model. Aberrant protein aggregation is sensitive to environmental and other factors, which determine the outcome of the aggregates (Rangachari et al., 2018; Condello and Stoehr, 2018). Using the Aβ-fatty acid system as a model that is known to adopt an alternative off-pathway, here we have employed a game-theoretic framework on simplified, reduced order models. First, the results re-confirmed our previos observation that fatty acid concentrations modulate Aβ aggregation pathways (Rana et al., 2017). Second, we discovered that the adoption of on- or off-pthway aggregates tightly depends on a set of rate contant ratios, which in turn suggest the thermodynamic stablity (equilibirum constants) of the emerging aggregates. Moreover, the α parameters are sensitive to the pseudo-micelle concentrations, L, which hold the key in modulating the pathways. Second, the models also provide insights into the feasibility of bridging pathways as a function of emerging higher order aggregates. For example, the reduced order, six-species model predict four different scnearios or dominant pathways of reactions which are strongly dependent upon the bridge, while also suggesting that ∝_4_ is the key to formation of larger aggregates in off-pathway. Stability arguments also show the larger aggregates in this system to be more stable. The EKS simulations display a similar outcome; the simulations indicate that the larger the oligomer, the more significant the impact of that bridge upon formation of the respective fibril.

In our experiments, we note that the propensity to switch pathways is highest when the order of aggregate is the lowest (low molecular weight) and increasingly become weaker as we move toward higher order aggregates along either pathways. This is actually in agreement with our theoretical observations noting the fact that in experiments, high molecular weight species refer to fibrils while low molecular weight aggregates then refer to the range of oligomers taken up in EKS and ROM simulations. Perhaps profound insight is that the model will be able to predict the emergence of certain oligomers by a set of kinetic and thermodynamic values expressed as afunction of *α* and *λ*, from a “win” or “lose” perspective (Figures 4 and 6).

A key observation of this paper emerges from Figure 6, which indicates the presence of multiple (neutrally) stable pathways, beyond simply the on and off-pathways. The hybrid pathways, especially the off-on domain shown in yellow in Figure 6, provides a range of possibilites for ∝_3_ and ∝_4_ to draw the aggregation dynamics away from toxicity. This is a particularly significant result suggesting possible intervention strategies. Numerical simualtions and experiments both clearly support this qualitative result (Figures 7 and 8), by revealing dilution to be a clear strategy to force the off-on transition. Although not in detail, the experiments do indicate that it is possible that via concentration-dependent introduction or removal of *L* to the reaction, one could populate oliomgers as predicted by ROM.

Results of ROM indicate that when the ratio of non-bridge forward to backward rates is close to unity, the model achieves equilibrium very quickly. However, when forward rates are considerably higher than backward rates, as is to be expected under normal circumstances, the system takes considerably longer to achieve equilibrium due to a cycle of over-shooting of species sizes resulting from a large difference in reaction rates. Similarly, corresponding EKS simulations indicate a *10^18^* fold difference in the forward and backward switching rate constants (Table A1 in Appendix) pointing to potentially irreversible effects of switching oligomers between pathways although the system may take a longer time to achieve equilibrium; this observation however, pertains to our reaction system with fixed initial monomer concentration and is expected to show fluctuating dynamics by considering monomer or pseudo-micelle entry rates and stochastic effects of the switching of oligomers between the pathways.

## 5. Conclusions

The results presented here showcase the applicability of classical game theory on understanding amyloid aggregation pathways. This is significant because it provides an ability to predict the emergnce of aggregates along multiple pathways along a temporal and equilibrium landscape map. Such a map can be further refined to see how it evolves as a function of a given interacting partner of Aβ, such as fatty acid as demonstrated here. Perhaps the most significant impact of this work is that the prediction of the emergence of oligomers provides a handle for understanding the conditions at which toxic strains could be generated. This simplified model presented here can be fine-tuned into more sphosticated models by including more species along pathways, additional pathways and more interacting partners that can modeulate the pathway etc. In sum, the results presented here establishes a new paradigm in understanding the complex dynamics of Aβ aggreagation and provides impetus towards deciphering amyloid pathogenesis along with making therapeutic and diagnostic advances for such debilitating diseases in the future.

## Acknowledgements

The authors want to thank the National Science Foundation for their financial support; NSF CBET 1802793 (to VR), NSF CBET 1802588 (to PG), and NSF CBET 1802641 (to AV). The authors also thank the National Center for Research Resources [5P20RR01647-11] and the National Institute of General Medical Sciences [8 P20 GM103476-11] from the National Institutes of Health for funding through INBRE (to V.R.).

## APPENDIX

Differential equation of the species:

On-pathway species:

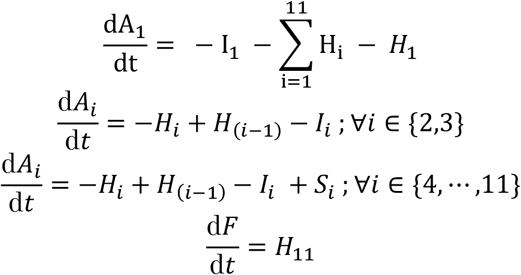

Off-pathway species:

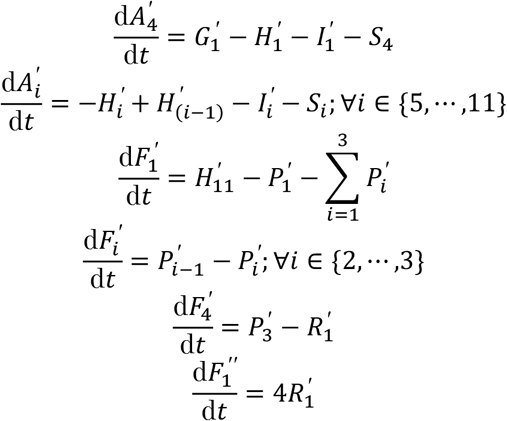

Pseudo-micelle

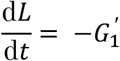

The simulated signal calculated as follow:

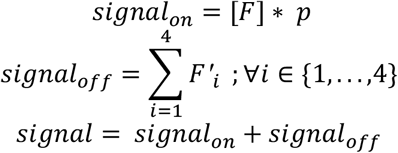

The estimated parameters are given below:

**Table A1:**
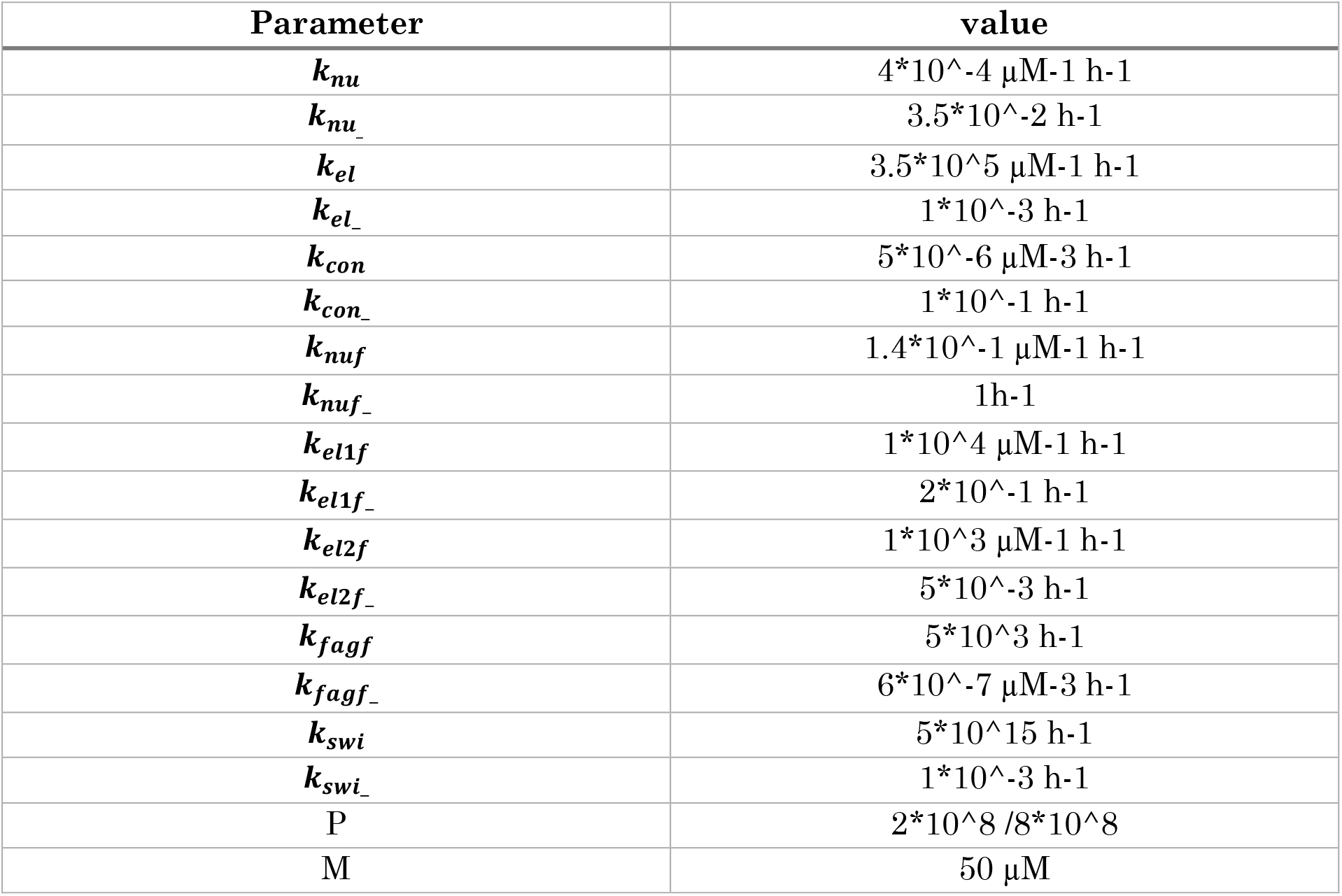
Estimated parameters from the EKS model.

| Parameter | value |
| --- | --- |
| $k_{nu}$ | $4 \cdot 10^{-4} \mu\text{M}^{-1} \text{h}^{-1}$ |
| $k_{nu\_}$ | $3.5 \cdot 10^{-2} \text{h}^{-1}$ |
| $k_{el}$ | $3.5 \cdot 10^5 \mu\text{M}^{-1} \text{h}^{-1}$ |
| $k_{el\_}$ | $1 \cdot 10^{-3} \text{h}^{-1}$ |
| $k_{con}$ | $5 \cdot 10^{-6} \mu\text{M}^{-3} \text{h}^{-1}$ |
| $k_{con\_}$ | $1 \cdot 10^{-1} \text{h}^{-1}$ |
| $k_{nuf}$ | $1.4 \cdot 10^{-1} \mu\text{M}^{-1} \text{h}^{-1}$ |
| $k_{nuf\_}$ | $1 \text{h}^{-1}$ |
| $k_{el1f}$ | $1 \cdot 10^4 \mu\text{M}^{-1} \text{h}^{-1}$ |
| $k_{el1f\_}$ | $2 \cdot 10^{-1} \text{h}^{-1}$ |
| $k_{el2f}$ | $1 \cdot 10^3 \mu\text{M}^{-1} \text{h}^{-1}$ |
| $k_{el2f\_}$ | $5 \cdot 10^{-3} \text{h}^{-1}$ |
| $k_{fagf}$ | $5 \cdot 10^3 \text{h}^{-1}$ |
| $k_{fagf\_}$ | $6 \cdot 10^{-7} \mu\text{M}^{-3} \text{h}^{-1}$ |
| $k_{swi}$ | $5 \cdot 10^{15} \text{h}^{-1}$ |
| $k_{swi\_}$ | $1 \cdot 10^{-3} \text{h}^{-1}$ |
| P | $2 \cdot 10^8 / 8 \cdot 10^8$ |
| M | $50 \mu\text{M}$ |

